# HOW FIVE DECADES OF LAND-COVER CHANGE RESHAPED SUITABLE HABITAT FOR PUERTO RICAN TREE SPECIES

**DOI:** 10.64898/2026.03.21.710527

**Authors:** Laura Moro, Pascal Milesi, Eileen Helmer, María Uriarte, Robert Muscarella

## Abstract

**Aim:** Human land-use has dramatically altered the amount, quality, and connectivity of habitat for species worldwide. Understanding how these changes affect individual species is essential for predicting the overall consequences of land-use change for biodiversity.

**Location:** The Caribbean island of Puerto Rico. Forest cover on the island increased from about 18 to 45% from the late 1940s to the early 2000s.

**Methods:** Using data on geographic distributions and functional traits for 454 tree species, we evaluated how gain of potential habitat was related to species-specific climatic associations and life-history strategies. We estimated species-specific potential habitat (climatically suitable and forested) with species distribution models and data on forest cover. We characterized each species’ niche breadth (the range of environmental conditions it occupies) and niche position (the environmental conditions it prefers) to compare with the conditions in reforested areas.

**Results:** Species with relatively more potential habitat in 1951 (climatically suitable and forested) also had relatively larger gains in potential habitat from 1951 to 2000. Species that tend to occupy conditions different from those common in reforested areas (i.e., more ‘marginal’ habitats) gained relatively less potential habitat and species with broad environmental niches gained more potential habitat. Additionally, species with relatively acquisitive functional traits gained more suitable habitat than those with relatively conservative traits.

**Main conclusions:** Our results show that Puerto Rico’s reforestation preferentially increased habitat for species that (1) already had suitable habitat in the landscape, (2) tolerate a wide range of climatic conditions, and (3) exhibit fast, acquisitive functional strategies. These findings illustrate how land-use change in heterogeneous tropical landscapes can generate non-uniform habitat gains across species, potentially favoring generalist over specialist species and reshaping community composition.

## INTRODUCTION

Human land-use can dramatically impact biodiversity by affecting the quality, amount, and connectivity of habitat (Fahrig, 2017; Hansen et al., 2012). While habitat loss and degradation due to land-use and land-cover change (LULCC) is recognized as a primary driver of biodiversity loss (IPBES, 2024; Magioli et al., 2021), land-use changes that lead to habitat gains are also widespread (Ellis et al., 2013; Kauppi et al., 2006; Mather, 1992; Rudel et al., 2009). For example, while the expansion of agricultural land remains a major driver of global forest cover loss (Diniz et al., 2022; Gibbs et al., 2010; Song et al., 2018), secondary forests now occupy large expanses of formerly agricultural land (Bousfield & Edwards, 2025; Chazdon et al., 2009, 2016). Understanding the consequences of LULCC in terms of both habitat loss and gain is thus crucial for assessment of biodiversity risk exposure and conservation planning.

A large body of research on the forest transition theory (Mather, 1992; Mather & Needle, 1998) has focused on the socio-economic and biophysical drivers of land-use change leading to forest regrowth. One key lesson from this work is that LULCC does not occur randomly across landscapes. Rather, areas that are more accessible and of higher value for human use (e.g., highly fertile) tend to be exploited first and abandoned last (Helmer et al., 2008; Huston, 2005; Scott et al., 2001). Moreover, protected areas (e.g., nature reserves) tend to be located in relatively less productive areas (Scott et al., 2001). The consequences of non-random patterns of land-use for biodiversity thus depend, in part, on how spatial patterns of land exploitation correspond to the particular environmental associations of individual species. Because species differ in their associations with different biophysical conditions, we may expect species-specific effects in terms of shifts in the amount and configuration of available habitat following LULCC (Hansen et al., 2012). Prior work on the consequences of LULCC for forest ecosystems has, however, focused largely on structural metrics (e.g., biomass), aggregate diversity patterns (e.g., species richness), species composition, or species functional groups (e.g., Fang et al., 2022; Gei et al., 2018; Helmer et al., 2018; Martinuzzi et al., 2022; Poorter et al., 2016). In comparison, we have a limited understanding of species-specific impacts of LULCC in different environments.

Characterizing species in terms of their niche position and niche breadth may help generalize the impact of LULCC across species (Figure 1) (Antão et al., 2022; Sexton et al., 2017). Niche position refers to the favored conditions for a species, and how those overlap with the conditions available in the landscape. Metrics of niche position can thus reflect the magnitude of difference between a species’ preferred conditions and the most commonly available conditions in a landscape (Morueta-Holme et al., 2013; Ohlemüller et al., 2008; Sheth et al., 2020). For example, species that prefer conditions similar to areas that become reforested would be expected to have larger gains in actual habitat compared to species that tend to occur in other conditions. Species niche breadth, or the range of conditions a species occupies, could also explain differences in the amount of available habitat following LULCC (Di Cecco & Hurlbert, 2022). Generalist species (i.e., those with a wider niche breadth) would typically be expected to experience larger gains of suitable habitat with LULCC compared to specialist species. Importantly, the niche position and niche breadth hypotheses are not mutually exclusive.

**Figure 1.**
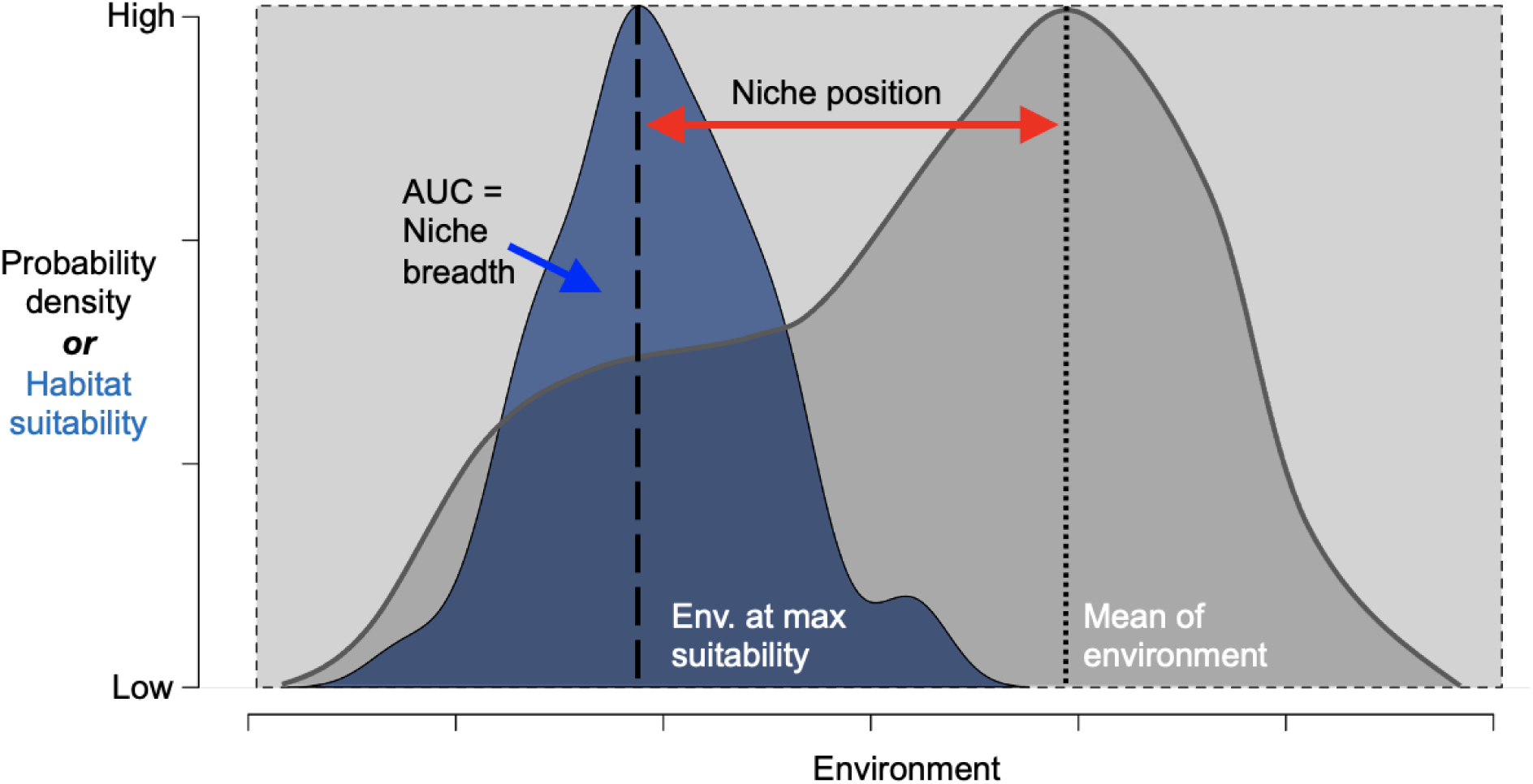
Schematic diagram illustrating niche position and niche breadth metrics for a hypothetical species. The grey curve represents the distribution of environmental values in the landscape. The blue curve represents the distribution of habitat suitability values (SDM predictions) along the same environmental gradient. Niche position is computed as the absolute difference between the mean value of environmental conditions in the landscape (dotted vertical line) and the environmental conditions of maximum habitat suitability for the species (dashed vertical line). Niche breadth is computed as the area under the habitat suitability curve, expressed as a proportion of the total area across the range of environment in the landscape (grey rectangle).

Traits that mediate life-history trade-offs (i.e., ‘functional traits’) may also help generalize species-specific responses in terms of habitat gains and losses following LULCC (Díaz, 2025). As the amount of potential habitat available across regional abiotic gradients changes, species traits influence differential species growth, survival, and recruitment across abiotic gradients could be related to patterns of habitat loss/gain. For example, trees in moist forests tend to have relatively low values of wood density (Muscarella & Uriarte, 2016). If LULCC predominantly occurs in moist forests, trees with traits more commonly associated with those regions should experience larger gains in habitat. However, forests also typically contain species with a large range of functional variation (Muscarella & Uriarte, 2016), which may indicate that traits are weak predictors of habitat gains/losses even if LULCC is biased with respect to abiotic conditions.

An ideal study system for assessing how niche position, niche breadth, and functional traits may relate to species-specific changes in potential habitat availability following LULCC would include substantial LULCC dynamics, high species diversity, and comprehensive data on geographic distributions and habitat preferences. The Caribbean island of Puerto Rico provides such a case study (Grau et al., 2003; Muscarella & Uriarte, 2016; Yackulic et al., 2011). While exact estimates vary, forest cover in Puerto Rico generally increased from about 18% of the total land area in 1950 to ca. 55% in 2022, following the widespread abandonment of agricultural land (Martinuzzi et al., 2022; Rudel et al., 2000). Prior work has related this large gain in forest cover to a variety of socio-economic and biophysical drivers (Grau et al., 2004; Kennaway & Helmer, 2007; Yackulic et al., 2011). Several studies have also investigated the implications of these changes for overall forest and land cover (Gould, 2009; Helmer et al., 2002; Lugo & Helmer, 2004), metrics of forest structure (e.g., biomass) (Martinuzzi et al., 2022), as well as the species and functional composition of forests (Brandeis et al., 2009; Muscarella et al., 2016). Although some species-specific habitat mapping has been done for Puerto Rican species (Gould et al., 2008), we lack a species-specific assessment of how potential habitat availability changed during this period.

Here we examined how LULCC in Puerto Rico during the second half of the 20th century (1950-2000) affected the amount of suitable habitat available for the majority (N=454) of the island’s tree species. We used a species-specific analysis of habitat suitability based on climatic and edaphic associations, and integrated data on species traits to further assess these changes. We also quantified the degree to which changes in suitable habitat affected fragmentation of habitat patches. We asked three main questions: (1) How did the amount and fragmentation (i.e., distance between patches and spatial aggregation) of potentially suitable habitat change for different Puerto Rican tree species from 1951 to 2000? (2) What is the range of variation of potential species richness in these reforested areas of Puerto Rico? (3) To what extent are species-specific shifts in suitable habitat availability related to species niche position, niche breadth, and functional traits?

We expect large differences among species in terms of habitat gain/loss. Since the largest bioclimatic zone in Puerto Rico is moist forest (Daly et al., 2003) and this is also where the largest gains of reforestation occurred (Helmer et al., 2002; Kennaway & Helmer, 2007), we expect the largest gains in suitable habitat to be for species that occur primarily in this forest type. We also expect relatively large gains of potentially suitable habitat for species with relatively wide niche breadths due to their ability to occupy in a wide range of conditions, as well as species relatively acquisitive functional traits.

## METHODS

### Study Area

The island of Puerto Rico (∼9,000 km^2^) encompasses six Holdridge life zones (Holdridge, 1947), with climate zones ranging from subtropical dry forest to subtropical lower montane rain forest (Brandeis et al., 2007). The island is characterized by steep environmental gradients with mean annual precipitation ranging from ca. 700–4,500 mm yr, and diverse geological substrates including volcanic, limestone, alluvial, and ultramafic bedrock (Bawiec, 1998). In the mid-1900s, forest cover on the island had been reduced to ∼18% due to land clearing for agriculture (T. Kennaway & Helmer, 2007), with some estimates as low as 6% (Birdsey & Weaver, 1987). A variety of political and socio-economic changes led to the widespread abandonment of agricultural land and the subsequent natural regeneration of forests (Aide et al., 1995; Brandeis et al., 2007; Grau et al., 2003; Yackulic et al., 2011). Forest cover on the island was estimated to have increased to ∼45% in the early 2000’s (Kennaway & Helmer, 2007), and to even as high as ∼55% as of 2022 (Martinuzzi et al., 2022).

### Habitat suitability models, niche position, and niche breadth

To characterize species-specific suitable habitat for each of 454 tree species, we built species distribution models (SDMs) with Maxent 3.4.1 (Phillips et al., 2017) using the R Package ENMeval v 2.0 (Kass et al., 2021; Muscarella, Galante, et al., 2014). We used a data set of 16,146 total occurrence records comprised of observations from the online databases GBIF (www.gbif.org), four herbaria (UPRRP, MAPR, NY, US), and georeferenced records from previously published studies, as well as our own observations. The mean (± sd) number of occurrence records per species was 40.6 (± 41.2). We used the ‘target-group’ background approach to account for spatial sampling bias in our occurrence dataset (Anderson, 2003). Specifically, when building an SDM for a given species, we used records of all other species as background points. We used four abiotic covariates in our SDMs: log-transformed precipitation in the driest month (mm), an index of precipitation seasonality (Feng et al., 2013), temperature in the hottest month (°C), and a categorical map of geological substrate (Bawiec, 1998). We downloaded temperature and precipitation data (climatological averages from 1963-1995) at 450 m resolution from the PRISM Climate Group (http://prism.oregonstate.edu).

We fit models using all pairwise combinations of five regularization multipliers (1-5) and four feature class combinations (“L”, “LQ”, “LQH”, “H”) and we evaluated the performance of SDMs using spatial cross-validation. For species with ≤15 occurrence records, we partitioned the data using the jackknife method (Hastie et al., 2009) while for species with >15 occurrences, we used the ‘checkerboard2’ method (Radosavljevic & Anderson, 2014) with an aggregation factor of 5. We selected the best models based on the lowest value of omission rate at the 10 percentile and then the highest value of AUC (Kass et al., 2020, 2021; Muscarella & Galante, 2014).

In downstream analyses, we primarily used continuous predicted values of SDMs. However, for analyses of habitat fragmentation and potential species richness (see next section), we converted continuous SDM predictions to binary maps of suitable vs. non-suitable habitat based on a species-specific threshold value that maximized the True Skill Statistic (TSS) (Allouche et al., 2006). We used the ‘ecospat.max.tss’ function of the ecospat v 4.4.1 R package (Broennimann et al., 2024) to compute the threshold TSS value for each species.

### Habitat amount and connectivity

We used maps of land cover from 1951 and 2000 (Helmer, Córdova, et al., 2023; Helmer et al., 2018; Kennaway & Helmer, 2007) to estimate the amount and configuration of climatically-suitable and available (i.e., forested) habitat through time for each species. We focus on this period because the vast majority of forest change occurred during these years, with forest cover remaining mostly stable from 2000 to the present (Marcano-Vega, 2023). Full details on land-cover classification maps can be found in (Helmer, Córdova, et al., 2023; Helmer et al., 2018; Kennaway & Helmer, 2007). Briefly, the forest cover map from 1951 (original scale 1:120,000 or ∼1,200 m) was based on aerial photos. The 2000 map was derived from Landsat TM/ETM images with 30 m pixels. The spatial resolution of the downloaded datasets was 30 m, whereas the climate data used for SDMs was ∼450 m. To match the spatial resolution of these datasets, we aggregated the landcover maps to match the coarser resolution climate data using the ‘resample’ function of the terra R package v. 1.8-70 (Hijmans, 2025).

To estimate changes in the amount of suitable habitat that was forested (i.e., ‘potential habitat’), we masked the suitability maps (i.e., SDM output for each species) with the forest cover map from 1951 and from 2000. Then, for each species, we summed values of predicted habitat suitability across all forested grid cells. We calculated the change in potential habitat amount as the difference of the total weighted potential forest habitat for each species from 1951 to 2000. Units of potential habitat change as presented are in map pixels weighted by modeled habitat suitability rather than units of area, per se.

To assess habitat fragmentation at each time point, we computed the mean of euclidean nearest-neighbor distance (km) between habitat patches in 1951 and 2000 for each species using the TSS thresholded maps and the ‘lsm_l_enn_mn’ function of the Landscape Metrics v. 2.0 R package (Hesselbarth et al., 2019). We chose this metric to reflect how the connectivity of potential habitat patches changed through the study period.

### Functional traits

We used data on five functional traits that represent life-history strategies expanded from previously published work (Muscarella & Uriarte, 2016). Specifically, we considered: wood density (WD g cm ^-3^), tree maximum height (m), leaf thickness (mm), leaf specific area (SLA g mm^2^ g^-1^), and seed dry mass (g). For seed dry mass, we obtained data from the Seed Information Database (SID) (https://ser-sid.org/) and Swenson and Umaña (2015). For other traits, we used measurements from 202 species across 12 forest plots representing the climatic and environmental gradient in Puerto Rico. This combined data set included a total of 1237 (54%) missing measurements across the 5 traits included. We used phylogenetic imputation to fill gaps in trait data with the R package Rphylopars (Goolsby et al., 2017) and using the phylogeny of Puerto Rican trees published by (Muscarella, Uriarte, et al., 2014). Prior to imputation the trait values were scaled and centered by subtracting the mean and dividing by standard deviation. After trait imputation, we performed a principal component analysis (PCA) including all traits, and we used the first principal component axis (PC1) as a multivariate trait to represent the acquisitive-conservative continuum of life history strategies.

### Statistical Analyses

We used habitat suitability maps (spatial predictions of SDMs) to compute metrics of niche position and breadth for each species (Figure 1). To do so, we first centered and scaled each environmental variable included in the SDMs across the island. Next, we computed the median value of each variable among grid cells that were classified as forest regrowth (i.e., non-forest in 1950 and forest in 2000). For each species, we computed the median value of each environmental variable across grid cells with the highest modeled suitability for that species. We then computed ‘niche position’ as the average difference between the median environmental values in reforested grid cells and in grid cells with highest suitability. To compute niche breadth, we binned each environmental variable into 100 bins and found the maximum predicted suitability across grid cells in each bin. We then computed the area under the curve and expressed this as a proportion of the potential area if a species had maximum habitat suitability across the full range of the environmental variable. In other words, a niche breadth value of 1 indicates that the species has its maximum predicted habitat suitability across all environmental conditions. Prior to this analysis, we centered and scaled values of niche position, niche breadth, and PC1 to facilitate direct comparison of the effect size of each term.

To answer our first question about how the amount and connectivity of suitable habitat changed for Puerto Rican tree species from 1951 to 2000, we compared the changes in potential habitat amount and connectivity with the respective values in 1951. To address our second question about potential species richness in reforested areas of Puerto Rico, we summed the thresholded SDM predictions across the island. Finally, we used multiple linear regression to answer our third question about the associations between changes in potential habitat (or connectivity) and niche breadth, niche position, and species traits:

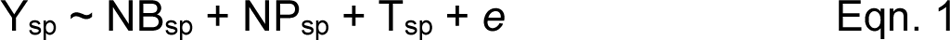

Where Y_sp_ is either the change in potential habitat or connectivity for species *sp*; NB is niche breadth, NP is niche position, T is the PCA axis 1 of functional traits, and *e* is a normally distributed error term. All statistical analyses were conducted in R version 4.5.2 (R Core Team, 2025).

## RESULTS

Based on maps from Helmer (2002) and Kennaway & Helmer (2007), Puerto Rico as a whole gained about 10 times as much forest as it lost between 1951 and 2000 (Figure 2). During this period, the largest gains of forest cover occurred in the two largest life zones on the island (subtropical moist and subtropical wet forest), which represent a total of 85% of the island’s land area (Table 1). Proportionally, the subtropical wet forest life zone gained the most forest cover with nearly 50% of its spatial extent transitioning to forest cover.

**Figure 2.**
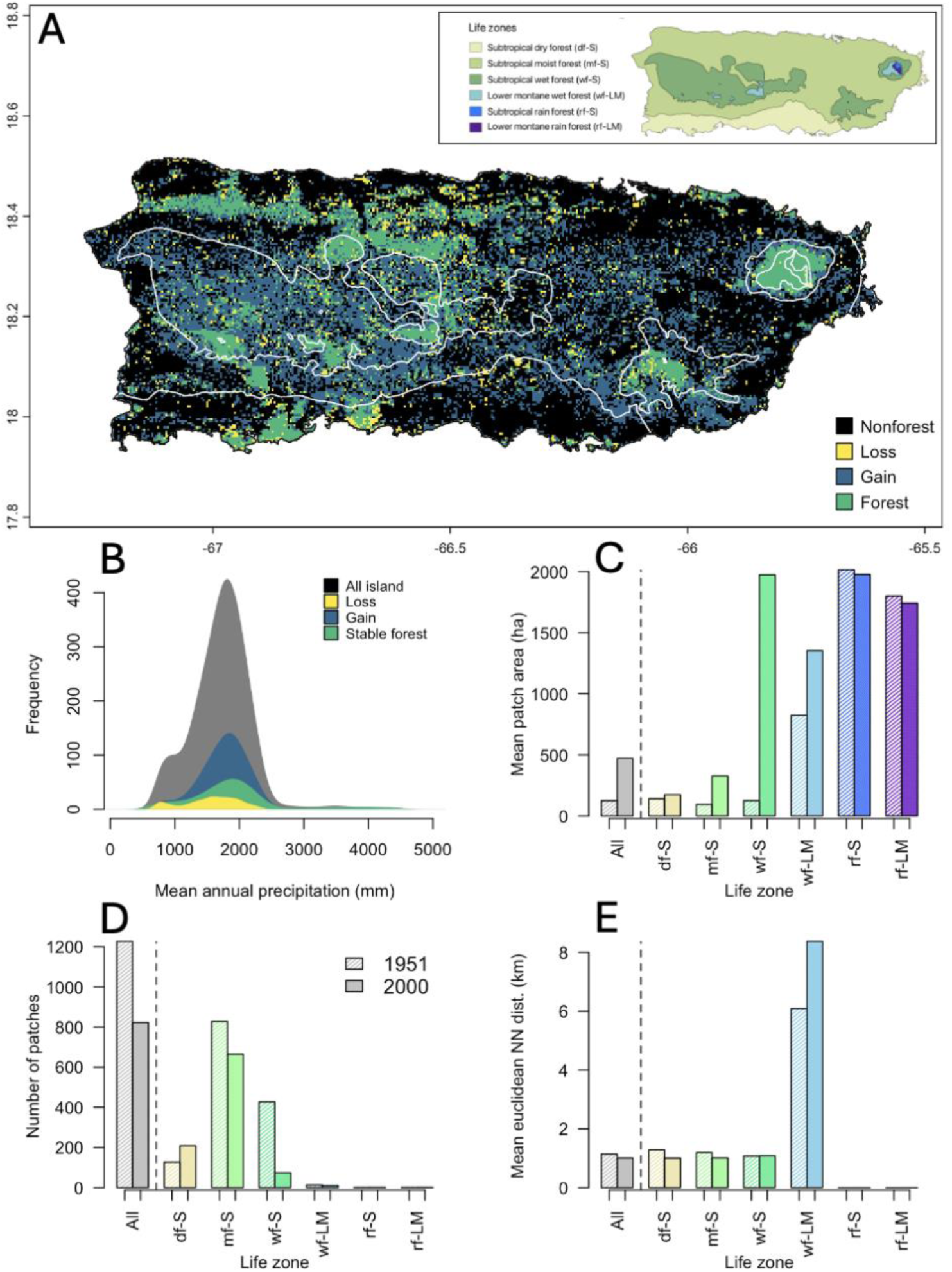
(A) Forest change map with persistent non-forest, forest lost from 1951-2000, forest gained from 1951-2000, persistent forest. White lines show delineations of life zones, as shown in the inset. (B) Density of grid cells along a gradient of mean annual precipitation for the different categories of LULCC in (A). (C-E) show the mean forest patch area (ha), total number of forest patches, and mean euclidean distance (km) between nearest neighbor forest patches across the entire island (grey bars) and within each life zone (colored bars) in 1951 (hatched bars) and in 2000 (solid bars). Bar colors in C-E correspond to the colors of panel (A) inset.

**Table 1.**
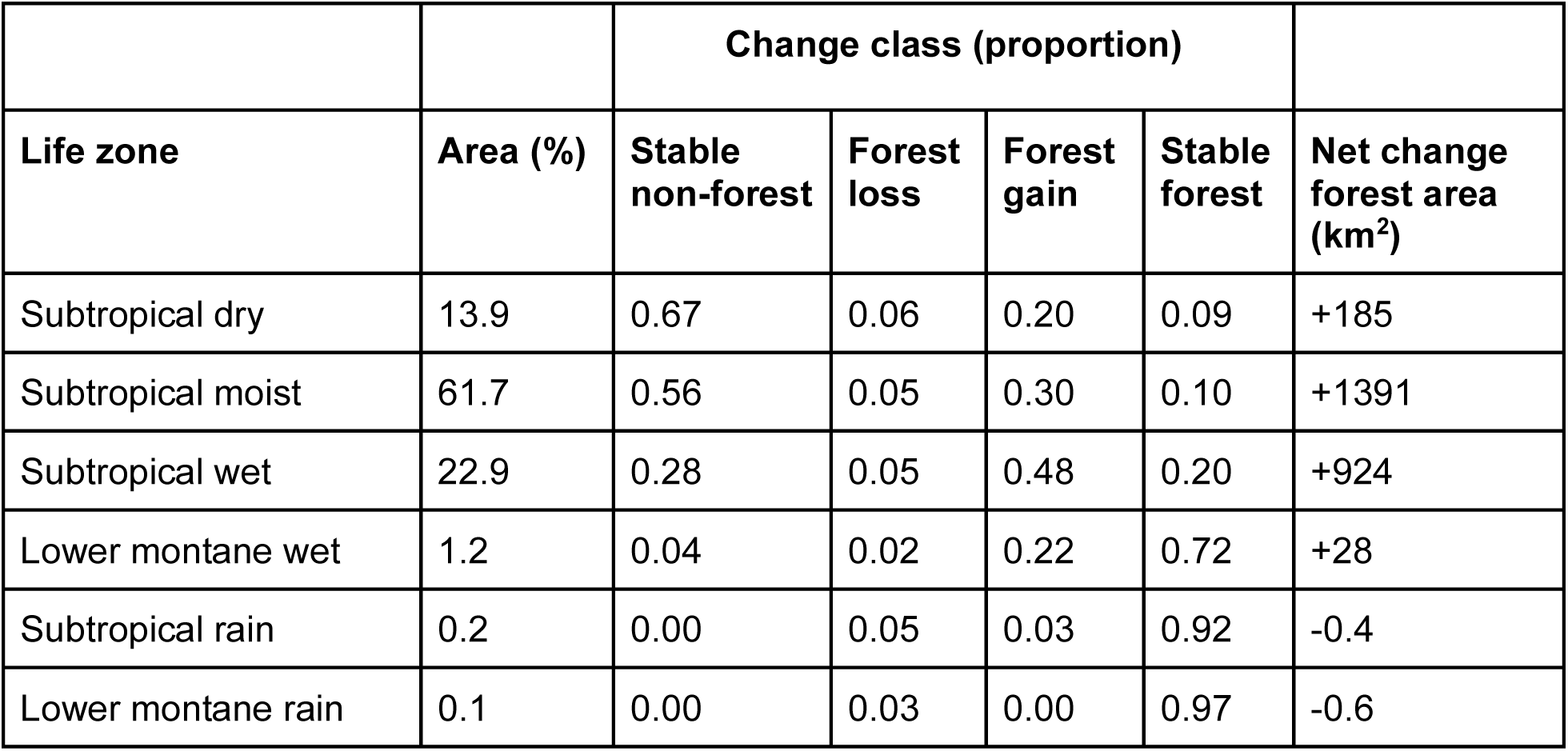
The percentage of the total area in mainland Puerto Rico classified into six different life zones (Daly et al., 2003), as well as the proportion of each life zone in different land cover change categories based on maps of land cover maps in 1951 and 2000 (Helmer, Córdova, et al., 2023; Helmer et al., 2002; Kennaway & Helmer, 2007).

Subtropical and lower montane rain forest (the two life zones with smallest total land area) had very little change in land cover with the vast majority remaining as stable forest. At the whole island scale, the average area of individual forest patches increased, while the total number of forest patches and nearest neighbor distance between patches decreased from 1951-2000 (Figure 2C-E).

The first two axes of the principal component analysis of traits explained 64% of the total variation (Figure 3A). In general, PC1 (which explained 42% of the total variation) was associated with the leaf economic spectrum; higher values represented relatively acquisitive traits (e.g., thin leaves and high SLA). PC2 (which explained an additional 22% of the variation) was most strongly related to wood density and leaf area; species with higher values of PC2 tended to have relatively large leaves and low density wood.

**Figure 3.**
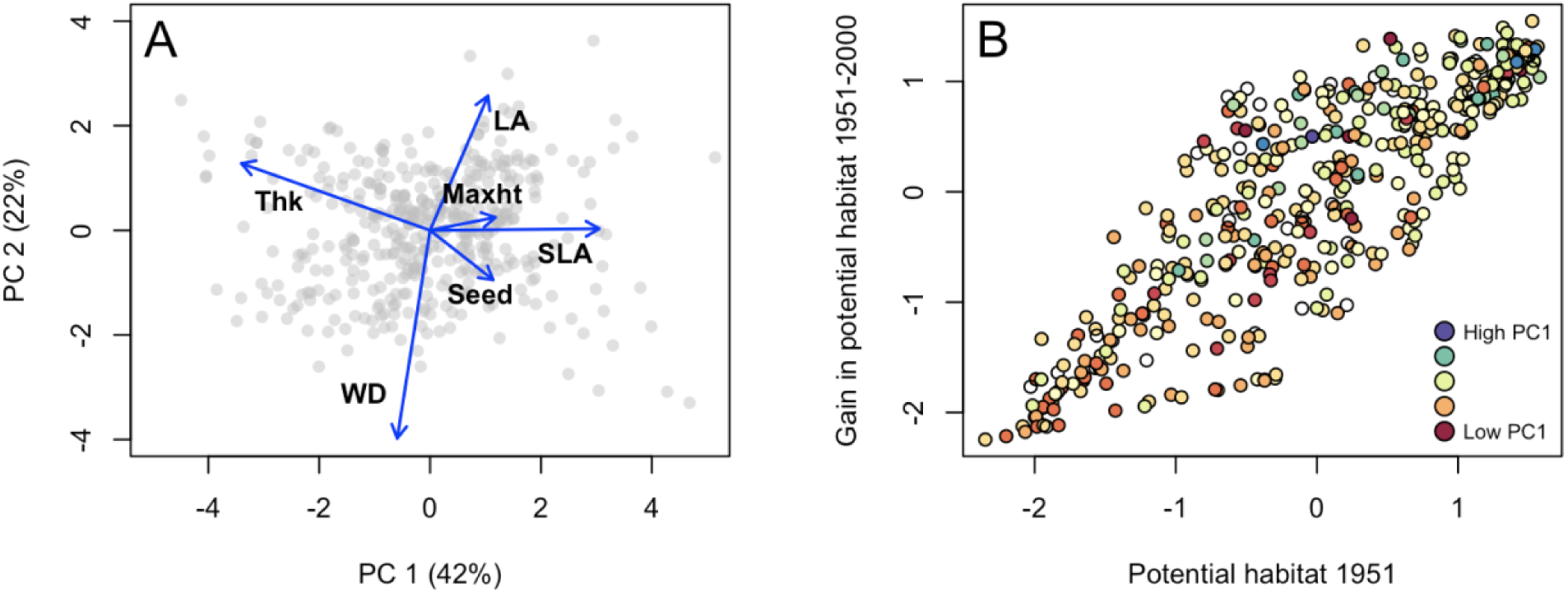
(A) Principal component axis 1 and 2 of the trait ordination analysis. Grey points are species scores, arrows show loadings of individual traits. The traits shown are wood density (WD, g cm ^-3^), tree maximum height (Maxht, m), leaf thickness (Thk, mm), leaf specific area (g SLA, mm^2^ g^-1^), and seed dry mass (Seed, g). (B) The amount (centered and scaled) of potential habitat per species in 1951 (x-axis) versus the amount (center and scaled) of increased potential habitat from 1951-2000. Each point represents one of 454 species and point colors show PC1 trait values.

Tree species with relatively more suitable and available habitat in 1951 also had relatively larger gains in potential suitable habitat from 1951 to 2000 (Figure 3B). Across species, the amount of potential suitable habitat gained via reforestation was 2–13 times more than the amount of suitable habitat lost (median ± SD = 6.4 ± 2.6). Across the island, reforested patches represented suitable habitat for 68-209 species (Figure 4). Predicted species richness in reforested patches was highest in the eastern parts of the Cordillera Central.

**Figure 4.**
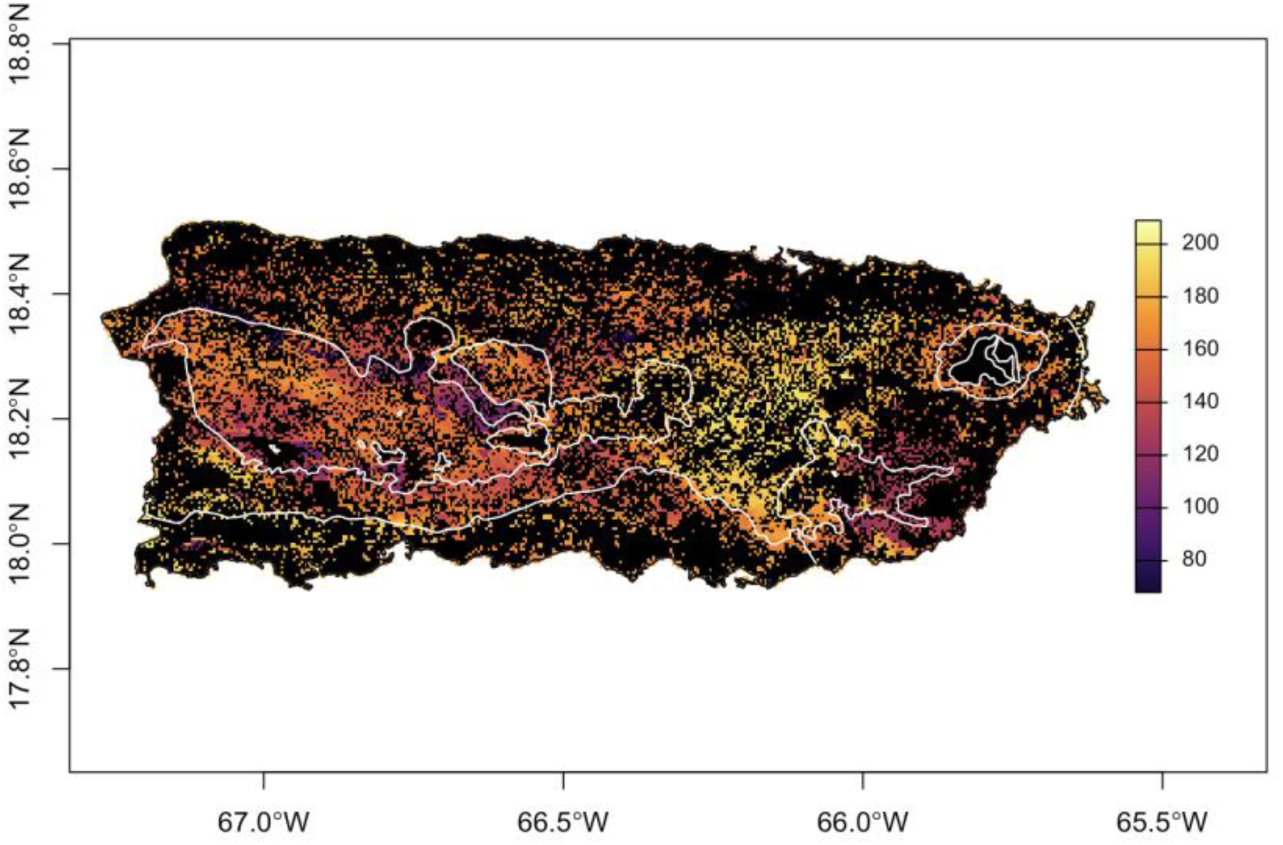
Potential species richness in forest regrowth from 1951 to 2000 based on overlapping maps of thresholded species distribution models and reforested pixels. White lines delineate life zones as in the inset of Figure 2A.

Metrics of niche position, niche breadth, and traits were weakly but significantly correlated across species. Niche breadth and niche position were negatively correlated (Pearson’s r = −0.36, p<0.001) indicating species that occupy more marginal habitats compared to the most common conditions in reforested areas also tend to have relatively smaller niche breadths. There was also a weak negative relationship between niche position and PC1 (Pearson’s r = −0.21, p<0.001), indicating that species with relatively acquisitive traits (higher values of PC1) tend to occur in relatively similar habitats compared to the conditions in reforested sites. There was a weak positive relationship (Pearson’s r = 0.31, p<0.001) between trait PC1 and niche breadth, indicating that species with more acquisitive traits tend to occur across a broader range of environmental conditions.

Niche position, niche breadth, and the PC1 trait axis were all significantly associated with the amount of potential habitat change from 1951-2000, with the combined linear model explaining 90% of the variation in the data (adjusted R^2^ = 0.90, Table S1). As expected, species that occupy more marginal habitats relative to the environmental conditions in the reforested areas (i.e., larger values of niche position) gained relatively less climatically-suitable habitat during the study period (p<0.001; Figure 5A). There was a stronger association between habitat gain and species niche breadth; species with broader niches had larger gains of suitable habitat (p<0.001; Figure 5B). Additionally, species with more acquisitive traits tended to have larger gains of suitable habitat, although the association was weaker than for either niche position or niche breadth (p=0.04; Figure 5C).

**Figure 5.**
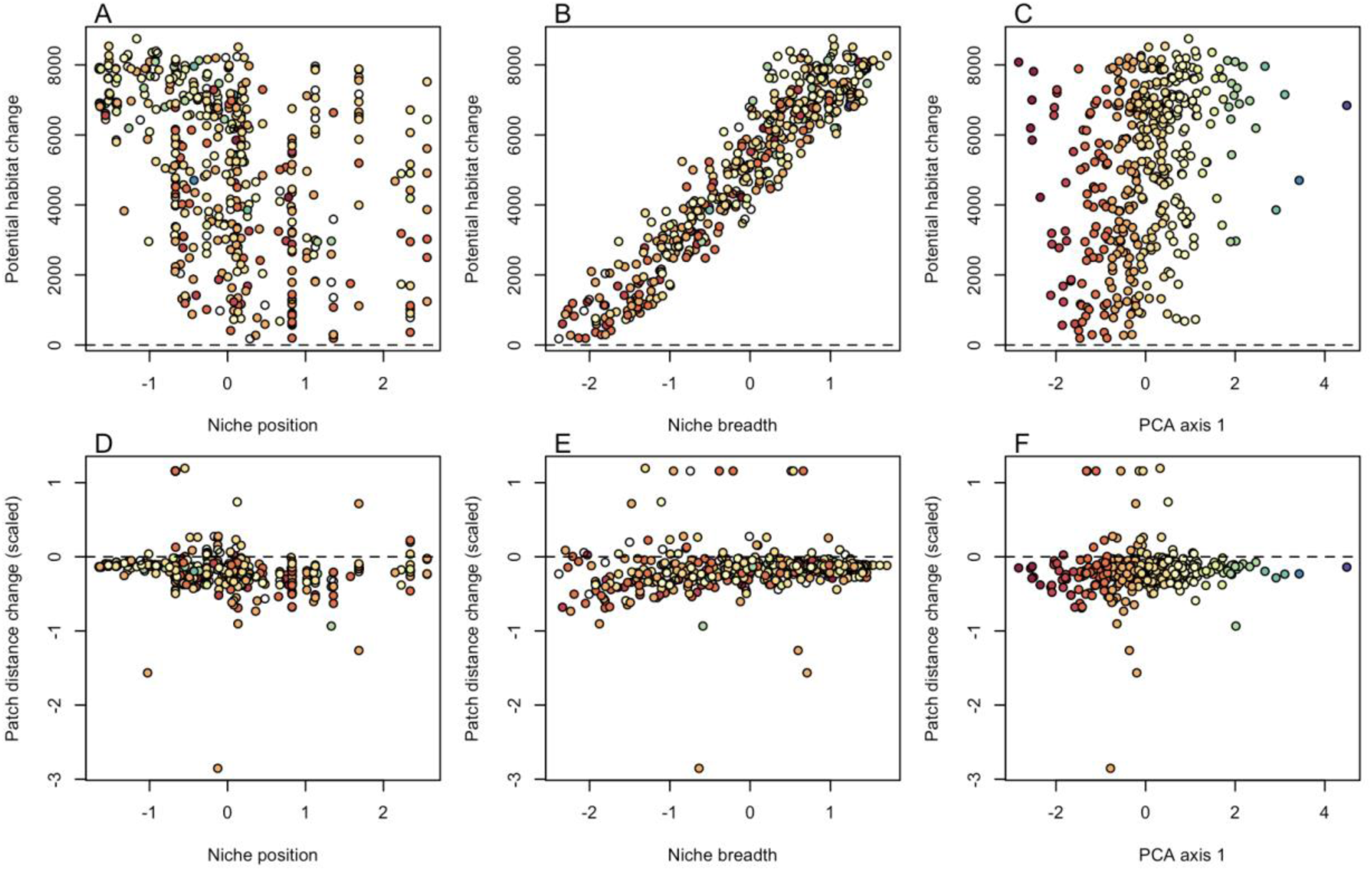
Relationship between the change in the amount (A-C) and mean distance between patches (D-F) of potential habitat from 1951 to 2000 as a function of (A, D) niche position (or marginality), (B, E) niche breadth, and (C, F) trait PCA for 454 Puerto Rican tree species. Point colors are based on values of trait PC1. Units for potential habitat change (A-C) are pixels weighted by habitat suitability; units for patch distance change (D-F) are scaled and centered kilometres. Point colors are based on values of trait PC1.

Reforestation also altered habitat connectivity – measured as nearest neighbor distance between potential habitat patches – differently across species but the combined model explained only ∼4% of the total variation (Figure 5D-F, Table S1). Species with lower values of niche position (i.e., those that tend to occur in areas with similar conditions as in reforested sites) had relatively large increases of connectivity (i.e., reduced distances between nearest neighbor patches of potential habitat) (p=0.02; Figure 5D). Species with lower values of niche breadth had relatively large decreases in terms of nearest neighbor distance between potential habitat patches (i.e., larger gains of potential connectivity) (p=0.01, Figure 5E). Change in connectivity was not significantly related to functional trait axis PC1 (p=0.97, Figure 5F).

## DISCUSSION

By the mid-1900s, Puerto Rico had been largely converted to agriculture, leaving limited habitat for forest-dwelling species, including trees. With the widespread abandonment of agricultural land during the second half of the 20th century, tree cover generally increased but spatial patterns of reforestation were non-random with respect to environmental conditions across the island, confirming prior studies (e.g., Helmer et al., 2008). Our study on the species-specific changes in availability of potential habitat from 1951 to 2000 showed: (1) The largest gains in potential habitat were realized by species with already large amounts of potential habitat in 1951; (2) Niche breadth (followed by niche position and species traits) was the strongest predictor for the amount of potential habitat gained during the study period, and (3) Niche breadth and niche position were weakly associated with changes in connectivity among patches (as measured by nearest neighbor distance). We discuss our results in terms of the consequences of natural forest regeneration for tropical tree biodiversity.

### Forest transitions

Global forest cover has expanded in recent decades, a trend that has been partly driven by agricultural abandonment (Rudel et al., 2009, 2020). In many instances, LULCC dynamics have occurred non-randomly across the landscape such that low-productivity and topographically rugged areas have been preferentially abandoned, leading to reforestation in these areas (Mather & Needle, 1998). In Puerto Rico, the forest transition happened mainly passively with selective abandonment of less-profitable lands in the steeper and wetter slopes of the central mountain, in rugged karst terrain and in other lands that are less arable or accessible (Helmer et al., 2008). In contrast, coastal plains remain largely used for agriculture and today are primary areas of urban development (Parés-Ramos et al., 2008). The spatial distribution of forests following these forest transitions are thus likely to systematically benefit some species over others, depending on environmental preferences.

Generally speaking, gains of forest habitat can carry a range of benefits for biodiversity and ecosystem services. For example, forest-dwelling species (including trees) may generally benefit from larger populations, which buffer against local extinction and can help safeguard genetic diversity (Chazdon et al., 2009). From an ecosystem services perspective, natural reforestation can provide enormous benefits (e.g., carbon sequestration) (Chazdon et al., 2016; Poorter et al., 2016). Numerous studies have shown, however, that species composition of regenerating secondary forests in diverse tropical regions is hardly predictable (Norden et al., 2015). The effects in terms of species-specific habitat recovery in the context of natural forest regeneration are often overlooked, yet they could provide important information to guide conservation practices and predict the potential for biodiversity recovery.

### Species-specific consequences of forest regeneration

In our study, species exhibited a wide range of potential habitat gains. The amount of habitat gain was most strongly correlated to species niche breadth, where broader niche species (habitat generalists) showed the largest gains of potential habitat. Niche position was also significantly associated with habitat gain whereby species that occupy more marginal habitats with respect to the most common conditions in the reforested landscape gained relatively less potential habitat. Notably, niche breadth and niche position were weakly (negatively) correlated across species such that species with central niche positions also tended to have high values of niche breadth. The largest increases in potential habitat gain between 1951 and 2000 were exhibited by species with central niche position and wide niche breadth. These results illustrate how natural forest regrowth can translate into relatively larger or smaller changes in potential habitat availability depending on species’ climatic associations.

In Puerto Rico, most of the land area falls within the subtropical moist and wet life zones, with comparatively little area in dry forest and the (wetter) higher elevation lower montane climatic zone. The largest areas of reforestation have also occurred in the dominant, moist and wet life zones. As a result, forest regrowth favored species whose niche positions more closely align with these dominant climatic conditions. Species with non-marginal niche positions tend to be more widespread and have higher occupancy rates and dispersal potential (Vela Díaz et al., 2020; Vleminckx et al., 2023). In our study system, species with niche positions closer to the conditions of the subtropical moist and wet life zones were more likely to become widespread during forest regeneration. As forest regrowth largely overlapped with their climatic optimum these species showed a greater gain in potential habitat.

It is perhaps not surprising that species with broader niche breadths exhibited the largest gains of potential habitat. This result suggests that habitat generalists are expected to benefit most during forest regeneration. A positive association between niche breadth and geographic range size is well established; generalist species tend to occupy larger ranges and often become ecologically dominant and locally abundant (Morueta-Holme et al., 2013; Moulatlet et al., 2025; Sheth et al., 2020). In Puerto Rico, we observe a similar pattern, with broad-niche species exhibiting substantially greater gains in potential habitat. As generalists, these species can occupy a wide range of environmental conditions, allowing most forest expansion to provide suitable habitat.

In our study, species’ traits were a significant (albeit relatively weak) predictor of potential habitat gain. The conservative–acquisitive plant economic spectrum, based on plant functional traits, is widely used to distinguish plant resource-use strategies (Wright et al., 2004) and can help predict recolonization and successional patterns after LULCC (Díaz, 2025; Poorter et al., 2021). In our study, species with conservative traits were associated with more marginal habitats and had narrower niche breadths (also see Helmer et al., 2018). A similar pattern was reported at the global scale by Chen et al. (2025), who found that species with conservative leaf traits were generally associated with narrow niche breadths. Overall we found that species with more acquisitive traits gained more potential habitat than species with more conservative traits.

Besides total forest area, forest cover increase in Puerto Rico also affected landscape configuration. Between 1951 and 2000 mean patch area increased and mean patch number and nearest neighbor distance between patches decreased. We interpret the change in these indices as a reduction in landscape fragmentation and an overall increase in landscape connectivity. A previous study examining habitat fragmentation changes in Puerto Rico between 1977 to 2000 found an increase in landscape aggregation of wetland habitats and a decrease in aggregation in forested habitats (Gao & Yu, 2014). Specifically Gao and Yu (2014) found that deforestation between 1977 to 2000 was located in the forest interiors, while the reforested sites tend to be located along the edges of the forests.

We found only a weak relationship between niche position and increases in connectivity (reflected by changes in nearest-neighbour distance). Both species with marginal and non-marginal niche position, showed significant increases in connectivity, although the effect was stronger for species with non-marginal niche position. Numerous studies highlight the crucial role of seed arrival in natural forest regeneration, suggesting that decreasing the distance between forest patches will facilitate dispersal and strongly influence the species composition of regenerating forest communities (Chazdon & Guariguata, 2016; Reid et al., 2015). When forest regrowth provides an increase in the suitable habitat available, its spatial configuration becomes an important determinant of habitat availability. Conversely, if regrowth does not provide an increase in suitable habitat, its extent may matter less, though it can still facilitate movement across the landscape (Bowen et al., 2007; Fricke et al., 2025). Percolation theory predicts that once a habitat covers more than about 59% of a landscape, connectivity does not limit species dispersal (Keitt et al., 1997). The ∼50-55% forest cover across Puerto Rico circa the year 2000 may mean that most habitat patches are relatively well-connected.

Maps of potential species richness are commonly generated in macroecological studies and can be valuable for applied research, conservation, and addressing fundamental questions about the processes governing spatial patterns of biodiversity (Conroy & Noon, 1996; Graham & Hijmans, 2006). We found a high degree of variation in potential species richness across reforested patches in Puerto Rico, ranging from 68 to 209 species per grid cell. This shows how reforestation has different potential to support biodiversity depending on where it takes place and which species are the main actors of the forest regrowth.

It is critical to note that our study focuses on quantifying the amount and arrangement of ‘potential’ habitat patches, as opposed to actually occupied habitat. This distinction is essential for considering the diverse consequences of LULCC for biodiversity and conservation. Individual species may be absent from high-quality patches for numerous reasons including, for example, dispersal limitation or priority effects. Additionally, the degree to which patches of climatically-suitable habitat are actually occupied by individual species also depends on highly local and dynamic conditions (e.g., understory light environment related to forest successional stage) that are not explicitly included in our distribution models. Thus, our estimates of potential suitable habitat could be further refined by considering how age (or age distribution) of suitability forest patches relates to successional associations for individual species. Moreover, tracking actual occupancy of forest patches through time across the island would provide additional insight to effects of LULCC on forest biodiversity. For example, a common, introduced, acquisitive species has relatively large island-wide basal area across Puerto Rico right after hurricane disturbances but declines as succession proceeds between them. For some groups of less acquisitive, native species, the basal area tended to have the opposite pattern or tended to increase over time (Helmer, Kay, et al., 2023b).

Natural forest regeneration has enormous potential to support biodiversity. Investigating the species-specific consequences of natural regeneration, and how it affects changes in potential habitat amount and configuration, can help predict and understand the trajectories of natural forest recovery and inform species conservation practices (Smith & Levine, 2025). In Puerto Rico, even though forest regrowth resulted in an overall increase in amount and connectivity of potential habitat for all tree species, the magnitude of potential habitat gain varied widely. In the context of LULCC and natural forest regeneration, some species tend to consistently be the “winners” in terms of habitat gain. This could become even more evident when examining the combined effects of climate change and LULCC. Sweeney and Jarzyna (2022) found that generalists (species with broader niche breadths) are more resilient to environmental change and benefit from natural regeneration at the expense of specialists (species with narrower niche breadths). These findings indicate that forests regenerating after clearing or other disturbances may have a community that is shifted toward acquisitive and generalist species, highlighting the importance of monitoring and supporting specialist and conservative species as forests recover.

## Conclusions

In Puerto Rico, the increase in forest cover over a 50 year period led to a large increase of potential habitat for tree species. The largest gains of potential habitat were realized by species that occupy similar habitats as those in the conditions of the reforested areas, species with relatively wide niche breadth, and species with relatively acquisitive functional traits. Natural forest regrowth in the heterogeneous landscapes of Puerto Rico led to complex patterns of habitat change across a diverse assemblage of species, which can have consequences for the diversity and composition of regenerating forests.

## Supporting information

Data for the manuscript

## ACKNOWLEDGEMENTS

Research was funded by grants from the Swedish Research Council, Formas (2020-00921 to RM and PM), the Swedish Research Council, Vetenskapsrådet (2019-03758 to RM), the Swedish Phytogeographical Society (to LM and RM), the Birgitta Sintring Foundation (to LM), and the US National Science Foundation (NSF DEB-1753810 to MU and RM). Additional support was provided by the Luquillo LTER (US NSF grant DEB-1831952), the University of Puerto Rico-Río Piedras, and the USDA Forest Service International Institute of Tropical Forestry.

## BIOSKETCH

Laura Moro is a doctoral student at Uppsala University. Her PhD thesis focuses on the effects of land-use dynamics on the geographic distributions, population sizes, and genetic diversity of tree species in Puerto Rico.

## DATA ACCESSIBILITY STATEMENT

Land cover maps for Puerto Rico in 1951 and 2000 can be obtained from Helmer *et al*. (2023a) and Kennaway and Helmer(2023). Climate data can be obtained from: https://prism.oregonstate.edu/. Summary data to reproduce analyses are provided in the supplemental materials.

## DISCLAIMERS

All opinions expressed in this paper are the authors’ and do not necessarily reflect the policies and views of the U.S. Department

## SUPPLEMENTAL MATERIALS

**Figure S1.**
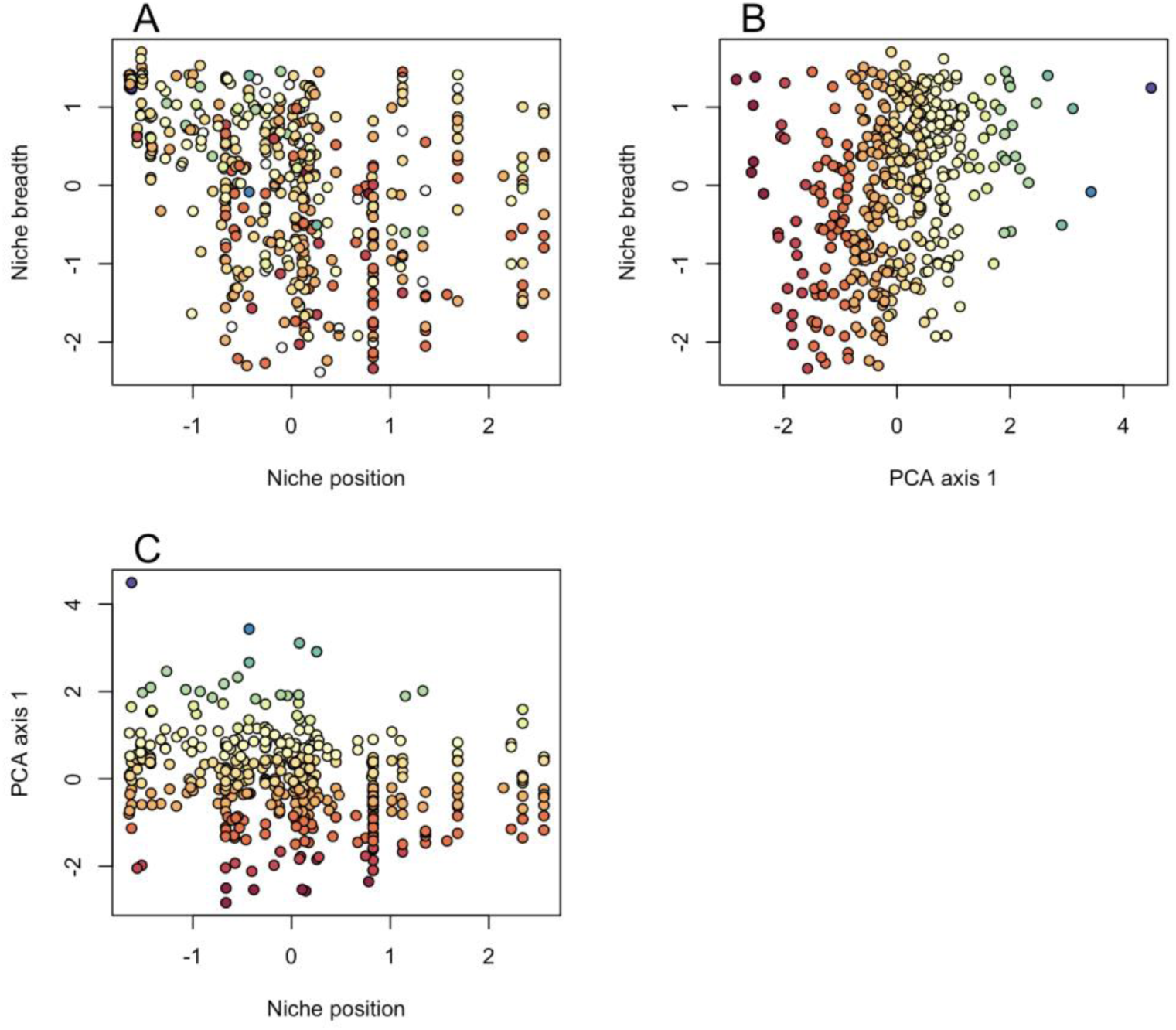
Correlations between niche breadth, niche position, and functional trait PCA axis 1.

**Table S1.**
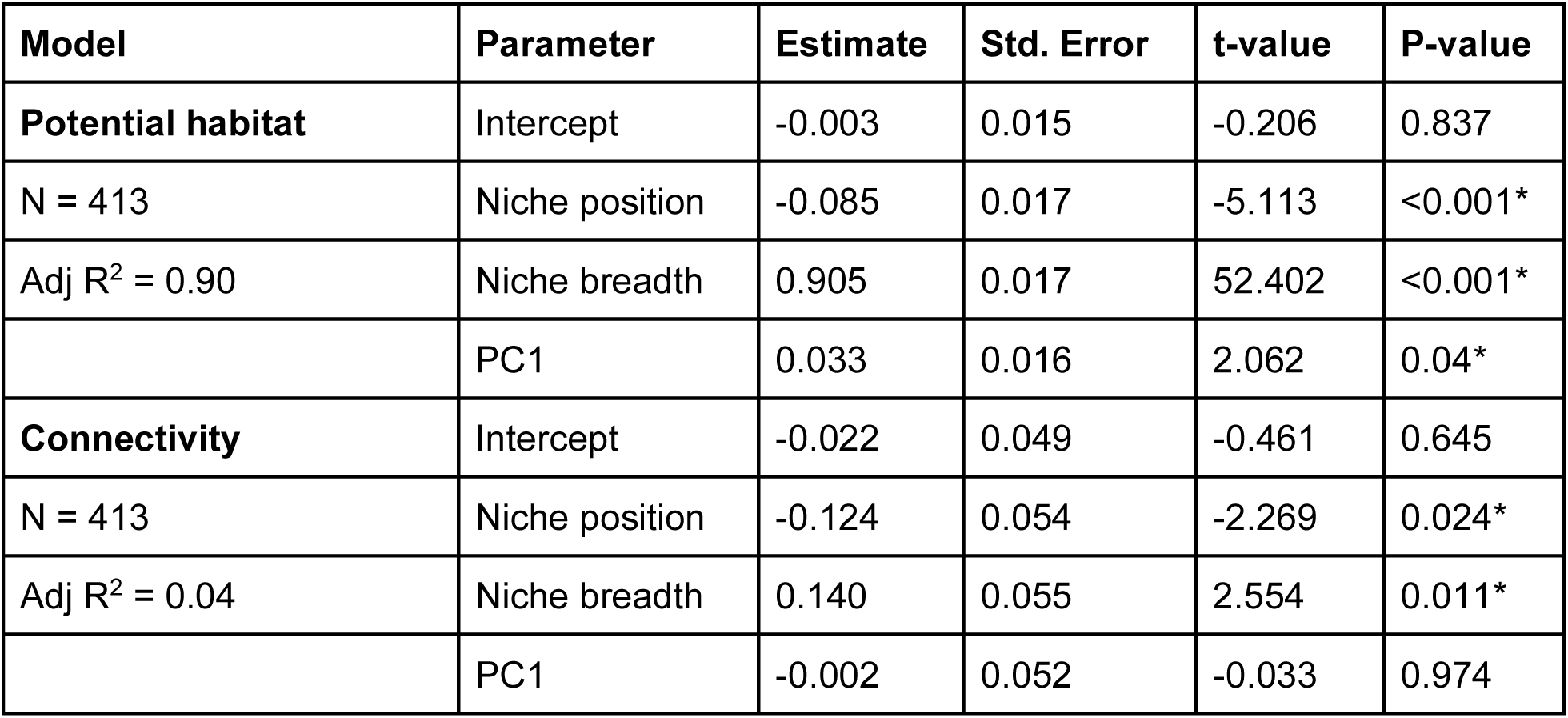
Summary statistics from multiple linear regression models.

## REFERENCES

Aide, T. M., Zimmerman, J. K., Herrera, L., Rosario, M., & Serrano, M. (1995). Forest recovery in abandoned tropical pastures in Puerto Rico. Forest Ecology and Management, 77(1), 77–86.

Allouche, O., Tsoar, A., & Kadmon, R. (2006). Assessing the accuracy of species distribution models: prevalence, kappa and the true skill statistic (TSS). The Journal of Applied Ecology, 43(6), 1223–1232.

Anderson, R. P. (2003). Real vs. artefactual absences in species distributions: tests for Oryzomys albigularis (Rodentia: Muridae) in Venezuela. Journal of Biogeography, 30(4), 591–605.

Antão, L. H., Weigel, B., Strona, G., Hällfors, M., Kaarlejärvi, E., Dallas, T., Opedal, Ø. H., Heliölä, J., Henttonen, H., Huitu, O., Korpimäki, E., Kuussaari, M., Lehikoinen, A., Leinonen, R., Lindén, A., Merilä, P., Pietiäinen, H., Pöyry, J., Salemaa, M., … Laine, A.-L. (2022). Climate change reshuffles northern species within their niches. Nature Climate Change, 12(6), 587–592.

Bawiec, W. J. (1998). Geology, geochemistry, geophysics, mineral occurrences, and mineral resource assessment for the Commonwealth of Puerto Rico. 10.3133/ofr9838

Birdsey, R. A., & Weaver, P. L. (1987). Forest Area Trends in Puerto Rico.

Bousfield, C. G., & Edwards, D. P. (2025). The pan-tropical age distribution of regenerating tropical moist forest. Nature Ecology & Evolution, 9(7), 1205–1213.

Bowen, M. E., McAlpine, C. A., House, A. P. N., & Smith, G. C. (2007). Regrowth forests on abandoned agricultural land: A review of their habitat values for recovering forest fauna. Biological Conservation, 140(3), 273–296.

Brandeis, T. J., Helmer, E. H., Marcano-Vega, H., & Lugo, A. E. (2009). Climate shapes the novel plant communities that form after deforestation in Puerto Rico and the U.S. Virgin Islands. Forest Ecology and Management, 258(7), 1704–1718.

Brandeis, T. J., Helmer, E. H., & Oswalt, S. N. (2007). The status of Puerto Rico’s forests, 2003. 10.2737/SRS-RB-119

Broennimann, O., Di Cola, V., & Guisan, A. (2024). ecospat: Spatial Ecology Miscellaneous Methods. https://CRAN.R-project.org/package=ecospat

Chazdon, R. L., Broadbent, E. N., Rozendaal, D. M. A., Bongers, F., Zambrano, A. M. A., Aide, T. M., Balvanera, P., Becknell, J. M., Boukili, V., Brancalion, P. H. S., Craven, D., Almeida-Cortez, J. S., Cabral, G. A. L., de Jong, B., Denslow, J. S., Dent, D. H., DeWalt, S. J., Dupuy, J. M., Durán, S. M., … Poorter, L. (2016). Carbon sequestration potential of second-growth forest regeneration in the Latin American tropics. Science Advances, 2(5), e1501639.

Chazdon, R. L., & Guariguata, M. R. (2016). Natural regeneration as a tool for large-scale forest restoration in the tropics: prospects and challenges. Biotropica, 48(6), 716–730.

Chazdon, R. L., Peres, C. A., Dent, D., Sheil, D., Lugo, A. E., Lamb, D., Stork, N. E., & Miller, S. E. (2009). The potential for species conservation in tropical secondary forests. Conservation Biology: The Journal of the Society for Conservation Biology, 23(6), 1406–1417.

Chen, Y., Hautier, Y., Kowalchuk, G. A., & Barry, K. E. (2025). “slow-fast” plant trait spectra are associated with ecological niches across global climatic gradients. Global Ecology and Biogeography: A Journal of Macroecology, 34(9), e70115.

Conroy, M. J., & Noon, B. R. (1996). Mapping of Species Richness for Conservation of Biological Diversity: Conceptual and Methodological Issues. Ecological Applications, 6(3), 763–773.

Daly, C., Helmer, E. H., & Quiñones, M. (2003). Mapping the climate of Puerto Rico, Vieques and Culebra. International Journal of Climatology, 23, 1359–1381.

Díaz, S. (2025). Plant functional traits and the entangled phenotype. Functional Ecology. 10.1111/1365-2435.70017

Di Cecco, G. J., & Hurlbert, A. H. (2022). Multiple dimensions of niche specialization explain changes in species’ range area, occupancy, and population size. Frontiers in Ecology and Evolution, 10, 921480.

Diniz, M. F., Coelho, M. T. P., Sánchez-Cuervo, A. M., & Loyola, R. (2022). How 30 years of land-use changes have affected habitat suitability and connectivity for Atlantic Forest species. Biological Conservation, 274(109737), 109737.

Ellis, E. C., Kaplan, J. O., Fuller, D. Q., Vavrus, S., Klein Goldewijk, K., & Verburg, P. H. (2013). Used planet: a global history. Proceedings of the National Academy of Sciences of the United States of America, 110(20), 7978–7985.

Fahrig, L. (2017). Ecological responses to habitat fragmentation per Se. Annual Review of Ecology, Evolution, and Systematics, 48(1), 1–23.

Fang, Z., Ding, T., Chen, J., Xue, S., Zhou, Q., Wang, Y., Wang, Y., Huang, Z., & Yang, S. (2022). Impacts of land use/land cover changes on ecosystem services in ecologically fragile regions. The Science of the Total Environment, 831(154967), 154967.

Feng, X., Porporato, A., & Rodriguez-Iturbe, I. (2013). Changes in rainfall seasonality in the tropics. Nature Climate Change, 3(9), 811–815.

Fricke, E. C., Cook-Patton, S. C., Harvey, C. F., & Terrer, C. (2025). Seed dispersal disruption limits tropical forest regrowth. Proceedings of the National Academy of Sciences of the United States of America, 122(30), e2500951122.

Gao, Q., & Yu, M. (2014). Discerning fragmentation dynamics of tropical forest and wetland during reforestation, urban sprawl, and policy shifts. PloS One, 9(11), e113140.

Gei, M., Rozendaal, D. M. A., Poorter, L., Bongers, F., Sprent, J. I., Garner, M. D., Mitchell Aide, T., Andrade, J. L., Balvanera, P., Becknell, J. M., Brancalion, P. H. S., AL Cabral, G., César, R. G., Chazdon, R. L., Cole, R. J., Colletta, G. D., De Jong, B., Denslow, J. S., Dent, D. H., … Powers, J. S. (2018). Legume abundance along successional and rainfall gradients in Neotropical forests. Nature Ecology & Evolution, 2(7), 1104–1111.

Gibbs, H. K., Ruesch, A. S., Achard, F., Clayton, M. K., Holmgren, P., Ramankutty, N., & Foley, J. A. (2010). Tropical forests were the primary sources of new agricultural land in the 1980s and 1990s. Proceedings of the National Academy of Sciences of the United States of America, 107(38), 16732–16737.

Goolsby, E. W., Bruggeman, J., & Ané, C. (2017). Rphylopars: fast multivariate phylogenetic comparative methods for missing data and within-species variation. Methods in Ecology and Evolution, 8(1), 22–27.

Gould. (2009). Puerto Rico Gap Analysis Project. GAP Analysis Bulletin. 16: 71-79, 16, 71–79.

Gould, W. A., Alarcon, C., Fevold, B., Jimenez, M. E., Martinuzzi, S., Potts, G., Quinones, M., Solórzano, M., & Ventosa, E. (2008). The Puerto Rico Gap Analysis Project volume 1: land cover, vertebrate species distributions, and land stewardship. Gen. Tech. Rep. IITF-39.

Graham, C. H., & Hijmans, R. J. (2006). A comparison of methods for mapping species ranges and species richness: Mapping species ranges and species richness. Global Ecology and Biogeography: A Journal of Macroecology, 15(6), 578–587.

Grau, H. R., Aide, T. M., Zimmerman, J. K., & Thomlinson, J. R. (2004). Trends and scenarios of the carbon budget in postagricultural Puerto Rico (1936–2060). Global Change Biology, 10, 1163–1179.

Grau, H. R., Aide, T. M., Zimmerman, J. K., Thomlinson, J. R., Helmer, E., & Zou, X. (2003). The Ecological Consequences of Socioeconomic and Land-Use Changes in Postagriculture Puerto Rico. BioScience, 53(12), 1159–1168.

Hansen, A. J., DeFries, R. S., & Turner, W. (2012). Land use change and biodiversity: A synthesis of rates and consequences during the period of satellite imagery. In Land Change Science (pp. 277–299). Springer Netherlands.

Hastie, T., Tibshirani, R., & Friedman, J. H. (2009). The elements of statistical learning: data mining, inference, and prediction. 2nd edn., Springer-Verlag.

Helmer, Brandeis, Lugo, & Kennaway. (2008). Factors influencing spatial pattern in tropical forest clearance and stand age: Implications for carbon storage and species diversity. Journal of Geophysical Research, 113(G2). 10.1029/2007jg000568

Helmer, E. H., Córdova, J. R., Quiñones, M., & Hubing, N. (2023). Historical maps of land use in Puerto Rico in 1951 [Dataset]. 10.2737/RDS-2023-0041

Helmer, E. H., Kay, S. L., Marcano-Vega, H., Powers, J. S., Wood, T. E., Zhu, X., Gwenzi, D., & Ruzycki, T. S. (2023a). Forest age map, tree species traits and Landsat phenology metrics for Puerto Rico and the U.S. Virgin Islands [Dataset]. In Forest Service Research Data Archive. USDA Forest Service. 10.2737/rds-2023-0004

Helmer, E. H., Kay, S., Marcano-Vega, H., Powers, J. S., Wood, T. E., Zhu, X., Gwenzi, D., & Ruzycki, T. S. (2023b). Multiscale predictors of small tree survival across a heterogeneous tropical landscape. PloS One, 18(3), e0280322.

Helmer, López, Díaz, & Piedras. (2002). Mapping the Forest type and land cover of Puerto Rico, a component of the Caribbean biodiversity hotspot. Caribbean Journal of Science, Vol. 38, No. 3-4, 165–183,. https://research.fs.usda.gov/treesearch/30146

Helmer, Ruzycki, Wilson, Sherrill, Lefsky, Marcano-Vega, Brandeis, Erickson, & Ruefenacht. (2018). Tropical Deforestation and Recolonization by Exotic and Native Trees: Spatial Patterns of Tropical Forest Biomass, Functional Groups, and Species Counts and Links to Stand Age, Geoclimate, and Sustainability Goals. Remote Sensing, 10(11), 1724.

Hesselbarth, M. H. K., Sciaini, M., With, K. A., Wiegand, K., & Nowosad, J. (2019). landscapemetrics: an open-source R tool to calculate landscape metrics. In Ecography (Vol. 42, pp. 1648–1657).

Hijmans, R. J. (2025). terra: Spatial Data Analysis. 10.32614/CRAN.package.terra

Holdridge, L. R. (1947). Determination of World Plant Formations From Simple Climatic Data. Science, 105(2727), 367–368.

Huston, M. A. (2005). The three phases of land-use change: Implications for biodiversity. Ecological Applications: A Publication of the Ecological Society of America, 15(6), 1864–1878.

IPBES. (2024). Thematic Assessment Report on the Underlying Causes of Biodiversity Loss and the Determinants of Transformative Change and Options for Achieving the 2050 Vision for Biodiversity of the Intergovernmental Science-Policy Platform on Biodiversity and Ecosystem Services (Transformative Change Assessment) (K. O’Brien, L. Garibaldi, & Agrawal A (eds.)). IPBES Secretariat.

Kass, J. M., Anderson, R. P., Espinosa-Lucas, A., Juárez-Jaimes, V., Martínez-Salas, E., Botello, F., Tavera, G., Flores-Martínez, J. J., & Sánchez-Cordero, V. (2020). Biotic predictors with phenological information improve range estimates for migrating monarch butterflies in Mexico. Ecography, 43(3), 341–352.

Kass, J. M., Muscarella, R., Galante, P. J., Bohl, C. L., Pinilla-Buitrago, G. E., Boria, R. A., Soley-Guardia, M., & Anderson, R. P. (2021). ENMeval 2.0: Redesigned for customizable and reproducible modeling of species’ niches and distributions. Methods in Ecology and Evolution / British Ecological Society, 12(9), 1602–1608.

Kauppi, P. E., Ausubel, J. H., Fang, J., Mather, A. S., Sedjo, R. A., & Waggoner, P. E. (2006). Returning forests analyzed with the forest identity. Proceedings of the National Academy of Sciences of the United States of America, 103(46), 17574–17579.

Keitt, T., Urban, D., & Milne, B. (1997). Detecting Critical Scales in Fragmented Landscapes. Conservation Ecology, 1(1). 10.5751/ES-00015-010104

Kennaway, & Helmer. (2007). The forest types and ages cleared for land development in Puerto Rico. GIScience and Remote Sensing, 44(4), 356–382.

Kennaway, T. A., & Helmer, E. H. (2023). Puerto Rico, Vieques and Culebra land cover and forest formations circa 2000 [Dataset]. In Forest Service Research Data Archive. USDA Forest Service. 10.2737/rds-2023-0022

Kennaway, T., & Helmer, E. H. (2007). The Forest types and ages cleared for land development in Puerto Rico. GIScience and Remote Sensing, 44(4), 356–382.

Lugo, A. E., & Helmer, E. (2004). Emerging forests on abandoned land: Puerto Rico’s new forests. Forest Ecology and Management, 190(2-3), 145–161.

Magioli, M., de Ferraz, K. M. P. M. B., Chiarello, A. G., Galetti, M., Setz, E. Z. F., Paglia, A. P., Abrego, N., Ribeiro, M. C., & Ovaskainen, O. (2021). Land-use changes lead to functional loss of terrestrial mammals in a Neotropical rainforest. Perspectives in Ecology and Conservation, 19(2), 161–170.

Marcano-Vega, H. (2023). Puerto Rico Forests 2019 (Vol. 461). U.S. Department of Agriculture, Forest Service. 10.2737/fs-ru-461

Martinuzzi, S., Cook, B. D., Helmer, E. H., Keller, M., Locke, D. H., Marcano-Vega, H., Uriarte, M., & Morton, D. C. (2022). Patterns and controls on island-wide aboveground biomass accumulation in second-growth forests of Puerto Rico. Biotropica, 54(5), 1146–1159.

Mather. (1992). The Forest Transition. Area, 24(4), 367–379.

Mather, A., & Needle, C. (1998). The forest transition: a theoretical basis. Area, 30(2), 117–124.

Morueta-Holme, N., Enquist, B. J., McGill, B. J., Boyle, B., Jørgensen, P. M., Ott, J. E., Peet, R. K., Símová, I., Sloat, L. L., Thiers, B., Violle, C., Wiser, S. K., Dolins, S., Donoghue, J. C, 2nd., Kraft, N. J. B., Regetz, J., Schildhauer, M., Spencer, N., & Svenning, J.-C. (2013). Habitat area and climate stability determine geographical variation in plant species range sizes. Ecology Letters, 16(12), 1446–1454.

Moulatlet, G. M., Merow, C., Maitner, B., Boyle, B., Feng, X., Frazier, A. E., Hinojo-Hinojo, C., Newman, E. A., Roehrdanz, P. R., Song, L., Villalobos, F., Marquet, P. A., Svenning, J.-C., & Enquist, B. J. (2025). General laws of biodiversity: Climatic niches predict plant range size and ecological dominance globally. Proceedings of the National Academy of Sciences of the United States of America, 122(46), e2517585122.

Muscarella, R., & Galante, P. J. (2014). ENMeval: An R package for conducting spatially independent evaluations and estimating optimal model complexity for Maxent ecological niche models. Methods in Ecology and Evolution / British Ecological Society. https://besjournals.onlinelibrary.wiley.com/doi/abs/10.1111/2041-210X.12261

Muscarella, R., Galante, P. J., Soley-Guardia, M., Boria, R. A., Kass, J. M., Uriarte, M., & Anderson, R. P. (2014). ENMeval: An R package for conducting spatially independent evaluations and estimating optimal model complexity forMaxentecological niche models. Methods in Ecology and Evolution / British Ecological Society, 5(11), 1198–1205.

Muscarella, R., & Uriarte, M. (2016). Do community-weighted mean functional traits reflect optimal strategies? Proceedings. Biological Sciences / The Royal Society, 283(1827), 20152434.

Muscarella, R., Uriarte, M., Aide, T. M., Erickson, D. L., Forero-Montaña, J., Kress, W. J., Swenson, N. G., & Zimmerman, J. K. (2016). Functional convergence and phylogenetic divergence during secondary succession of subtropical wet forests in Puerto Rico. Journal of Vegetation Science: Official Organ of the International Association for Vegetation Science, 27(2), 283–294.

Muscarella, R., Uriarte, M., Erickson, D. L., Swenson, N. G., Zimmerman, J. K., & Kress, W. J. (2014). A well-resolved phylogeny of the trees of Puerto Rico based on DNA barcode sequence data. PloS One, 9(11), e112843.

Norden, N., Angarita, H. A., Bongers, F., Martínez-Ramos, M., Granzow-de la Cerda, I., van Breugel, M., Lebrija-Trejos, E., Meave, J. A., Vandermeer, J., Williamson, G. B., Finegan, B., Mesquita, R., & Chazdon, R. L. (2015). Successional dynamics in Neotropical forests are as uncertain as they are predictable. Proceedings of the National Academy of Sciences, 112(26), 8013–8018.

Ohlemüller, R., Anderson, B. J., Araújo, M. B., Butchart, S. H. M., Kudrna, O., Ridgely, R. S., & Thomas, C. D. (2008). The coincidence of climatic and species rarity: high risk to small-range species from climate change. Biology Letters, 4(5), 568–572.

Parés-Ramos, I., Gould, W., & Aide, T. (2008). Agricultural Abandonment, Suburban Growth, and Forest Expansion in Puerto Rico between 1991 and 2000. Ecology and Society, 13(2). 10.5751/ES-02479-130201

Phillips, S. J., Research, A., Anderson, R. P., Schapire, R. E., & Dudik, M. (2017). A Brief Tutorial on Maxent. https://biodiversityinformatics.amnh.org/open_source/maxent/Maxent_tutorial2017.pdf

Poorter, L., Bongers, F., Mitchell Aide, T., Almeyda Zambrano, A. M., Balvanera, P., Becknell, J. M., Boukili, V., Brancalion, P. H. S., Broadbent, E. N., Chazdon, R. L., Craven, D., de Almeida-Cortez, J. S., AL Cabral, G., De Jong, B. H. J., Denslow, J. S., Dent, D. H., DeWalt, S. J., Dupuy, J. M., Durán, S. M., … Rozendaal, D. M. A. (2016). Biomass resilience of Neotropical secondary forests. Nature, 530(7589), 211–214.

Poorter, L., Rozendaal, D. M. A., Bongers, F., de Almeida, J. S., Álvarez, F. S., Andrade, J. L., Arreola Villa, L. F., Becknell, J. M., Bhaskar, R., Boukili, V., Brancalion, P. H. S., César, R. G., Chave, J., Chazdon, R. L., Dalla Colletta, G., Craven, D., de Jong, B. H. J., Denslow, J. S., Dent, D. H., … Westoby, M. (2021). Functional recovery of secondary tropical forests. Proceedings of the National Academy of Sciences of the United States of America, 118(49). 10.1073/pnas.2003405118

Radosavljevic, A., & Anderson, R. P. (2014). Making better Maxent models of species distributions: complexity, overfitting and evaluation. Journal of Biogeography, 41(4), 629–643.

R Core Team. (2025). R: A Language and Environment for Statistical Computing. R Foundation for Statistical Computing. https://www.R-project.org/

Reid, J. L., Holl, K. D., & Zahawi, R. A. (2015). Seed dispersal limitations shift over time in tropical forest restoration. Ecological Applications: A Publication of the Ecological Society of America, 25(4), 1072–1082.

Rudel, T. K., Meyfroidt, P., Chazdon, R., Bongers, F., Sloan, S., Grau, H. R., Van Holt, T., & Schneider, L. (2020). Whither the forest transition? Climate change, policy responses, and redistributed forests in the twenty-first century. Ambio, 49(1), 74–84.

Rudel, T. K., Perez-Lugo, M., & Zichal, H. (2000). When fields revert to forest: development and spontaneous reforestation in post-war Puerto Rico. The Professional Geographer: The Journal of the Association of American Geographers, 52, 386–397.

Rudel, T. K., Schneider, L., Uriarte, M., Turner, B. L., DeFries, R., Lawrence, D., Geoghegan, J., Hecht, S., Ickowitz, A., Lambin, E. F., Birkenholtz, T., Baptista, S., & Grau, R. (2009). Agricultural intensification and changes in cultivated areas, 1970–2005. Proceedings of the National Academy of Sciences, 106(49), 20675.

Scott, M. J., Davis, F. W., Gavin McGhie, R., Gerald Wright, R., Groves, C., & Estes, J. (2001). Nature reserves: do they capture the full range of America’s biological diversity? Ecological Applications, 11(4), 999–1007.

Sexton, J. P., Montiel, J., Shay, J. E., Stephens, M. R., & Slatyer, R. A. (2017). Evolution of ecological niche breadth. Annual Review of Ecology, Evolution, and Systematics, 48(1), 183–206.

Sheth, S. N., Morueta-Holme, N., & Angert, A. L. (2020). Determinants of geographic range size in plants. The New Phytologist, 226(3), 650–665.

Smith, J. R., & Levine, J. M. (2025). Linking relative suitability to probability of occurrence in presence-only species distribution models: Implications for global change projections. Methods in Ecology and Evolution, 16(4), 854–865.

Song, X.-P., Hansen, M. C., Stehman, S. V., Potapov, P. V., Tyukavina, A., Vermote, E. F., & Townshend, J. R. (2018). Global land change from 1982 to 2016. Nature, 560(7720), 639–643.

Sweeney, C. P., & Jarzyna, M. A. (2022). Assessing the synergistic effects of land use and climate change on terrestrial biodiversity: Are generalists always the winners? Current Landscape Ecology Reports, 7(4), 41–48.

Swenson, N. G., & Umana, M. N. (2015). Data from: Interspecific functional convergence and divergence and intraspecific negative density dependence underlie the seed-to-seedling transition in tropical trees [Dataset] [Dataset]. 10.5061/dryad.j2r53

Vela Díaz, D. M., Blundo, C., Cayola, L., Fuentes, A. F., Malizia, L. R., & Myers, J. A. (2020). Untangling the importance of niche breadth and niche position as drivers of tree species abundance and occupancy across biogeographic regions. Global Ecology and Biogeography: A Journal of Macroecology, 29(9), 1542–1553.

Vleminckx, J., Barrantes, O. V., Fortunel, C., Paine, C. E. T., Bauman, D., Engel, J., Petronelli, P., Dávila, N., Rios, M., Valderrama Sandoval, E. H., Mesones, I., Allié, E., Goret, J.-Y., Draper, F. C., Guevara Andino, J. E., Béroujon, S., Fine, P. V. A., & Baraloto, C. (2023). Niche breadth of Amazonian trees increases with niche optimum across broad edaphic gradients. Ecology, 104(7), e4053.

Wright, I. J., Reich, P. B., Westoby, M., Ackerly, D. D., Baruch, Z., Bongers, F., Cavender-Bares, J., Chapin, T., Cornelissen, J. H. C., Diemer, M., Flexas, J., Garnier, E., Groom, P. K., Gulias, J., Hikosaka, K., Lamont, B. B., Lee, T., Lee, W., Lusk, C., … Villar, R. (2004). The worldwide leaf economics spectrum. Nature, 428(6985), 821–827.

Yackulic, C. B., Fagan, M., Jain, M., Jina, A., Lim, Y., Marlier, M., Muscarella, R., Adame, P., DeFries, R., & Uriarte, M. (2011). Biophysical and socioeconomic factors associated with forest transitions at multiple spatial and temporal scales. Ecology and Society, 16(3). 10.5751/ES-04275-160315

